# Spatial coding in the hippocampus of flying owls

**DOI:** 10.1101/2021.10.24.465553

**Authors:** Arpit Agarwal, Ayelet Sarel, Dori Derdikman, Nachum Ulanovsky, Yoram Gutfreund

## Abstract

The elucidation of spatial coding in the hippocampus requires exploring diverse animal species. While robust place-cells are found in the mammalian hippocampus, much less is known about spatial coding in the hippocampus of birds – and nothing is known about avian spatial representation during flight. Here we used a wireless-electrophysiology system to record single neurons in the hippocampus and related pallial structures from freely flying barn owls (*Tyto alba*) – a central-place nocturnal predator species with excellent navigational abilities. The owl’s 3D position was monitored while it flew back and forth between two perches. We found place cells – neurons that robustly represented the owl’s location during flight, and its flight-direction – as well as neurons that coded the owl’s perching position between flights. Spatial coding was invariant to changes in lighting conditions and to the position of a salient object in the room. Place cells were found in the anterior hippocampus and in the adjacent posterior hyperpallium apicale, and to a much lesser extent in the visual Wulst (visual-cortex homologue). The finding of place-cells in flying owls suggests commonalities in spatial coding across a variety of species – including rodents, bats and owls.

## Introduction

A striking feature of the hippocampal formation of rodents and other mammalian species is the robust neural representation of space, e.g. place cells, grid cells and head-direction cells (Finkelstein et al., 2015; Hafting et al., 2005; O’Keefe and Dostrovsky, 1971; Taube et al., 1990; Yartsev et al., 2011). Such cells are thought to form the substrate of spatial memory and spatial perception of mammals (Eichenbaum, 2017; Moser et al., 2015; O’Keefe and Nadel, 1978). An important question is to what extent similar cells can be found in the brains of other, non-mammalian species – such as birds.

The avian hippocampal formation (Hp) lies in the posterior part of the forebrain, immediately below the dorsal surface of the brain (Atoji and Wild, 2006; Chen et al., 2013; Colombo and Broadbent, 2000; Herold et al., 2015; Reiner et al., 2005; Székely, 1999). Directly anterior to the Hp extends the hyperpallium, considered to be a homologue of the mammalian neocortex (Karten, 2015; Medina and Abellán, 2009; Stacho et al., 2020). However, the cytoarchitecture of the Hp and the hyperpallium is noticeably different from their mammalian homologues (Herold et al., 2015; Stacho et al., 2020). Lesion studies and immediate early gene activation suggest that the Hp and the hyperpallium play central roles in spatial cognition (Colombo and Broadbent, 2000; Gagliardo et al., 1999; Sherry et al., 2017; Smulders and DeVoogd, 2000; Watanabe and Bischof, 2004; Watanabe et al., 2011).

Reports on single-unit activity in the Hp of avian species portrayed diverse results: In quails, head-direction cells were found, but no place-cells (Ben-Yishay et al., 2021). In walking pigeons, studies reported multi-field neurons and cells that code the locations near rewards and/or the direction towards rewards (Bingman et al., 2003; Hough and Bingman, 2004; Kahn and Bingman, 2004; Kahn et al., 2008). In zebra finches, relatively few place-cells were found, restricted to the anterior part of the Hp (Payne et al., 2021). In a specialized food-caching bird (tufted titmouse) numerous and robust place cells were found during walking/hopping (Payne et al., 2021). The seemingly diverse spatial coding across avian species calls for further exploration. In particular, we hypothesized that robust place-cells would be found in birds with outstanding navigational skills and spatial memory capabilities, such as: (i) food-caching birds like titmouse and jays – which can memorize the exact locations of hundreds of cached food items (Grodzinski and Clayton, 2010); (ii) migrating species such as storks – which navigate thousands of kilometers to arrive at the exact same breeding location every year (Flack et al., 2018); or (iii) central-place foraging species – which excel at homing back to their preferred sites.

Here, we studied a novel avian model for the neurobiology of spatial memory – the barn owl. Barn owls are central-place foragers – nocturnal hunters which can self-localize themselves very well even in extremely low light levels, and navigate accurately back to their preferred perching-branches (Payne, 1971; Rozman et al., 2021). While barn owls were extensively studied in sensory neuroscience due to their excellent hearing and vision (Konishi, 2000; Pena and Gutfreund, 2014), their hippocampal formation was never studied electrophysiologically. We reasoned that due to their outstanding spatial orientation capabilities in darkness and strong central-place foraging behavior (Rozman et al., 2021), we would find in the owl’s hippocampus place-cells akin to those found in rodents, bats, and food-caching birds. Further, we hypothesized that place-cells would be found during flight – which was never tested to date in birds.

To this end, we implanted tetrode microdrives in several areas of the dorsal pallium – including in the hippocampal formation – and used a lightweight wireless-electrophysiology system to record single neurons from owls, as they were flying back and forth between two perches at opposite sides of a room. We found a plethora of spatially modulated neurons, including place cells: neurons that fired when the owl flew through a spatially restricted place-field in one direction but not in the other. These various spatially modulated cells were found in the anterior part of the hippocampus (Hp) and in the adjacent posterior part of the hyperpallium apicale (HA_p_), but to a lesser extent in the central part of the visual Wulst – a region considered to be the homologue of visual cortex (Pettigrew and Konishi, 1976). This study thus reveals a robust allocentric spatial representation in the owl’s brain, including spatially-localized place cells.

## Results

Previous electrophysiological and lesion studies in birds suggested that the anterior Hp is involved in spatial representation and spatial cognition (Lormant et al., 2020; Payne et al., 2021; Watanabe and Bischof, 2004). Therefore, in this study we searched for spatial coding in the anterior Hp and the adjacent HA_p_. To compare with results from a primary visual area, we also recorded in the visual Wulst, the owl’s homologue of the visual cortex (Pettigrew and Konishi, 1976). Overall, 761 well-isolated single units were recorded from six barn owls: 292 from Hp, 376 from HA_p_ and 93 from visual Wulst (see Table S1).

### Spatially modulated cells in the Hp

To target the anterior Hp, we directed the tetrodes 4 mm anterior to the cerebellum edge and 2 mm lateral from the midline (Fig. 1A, top). A reconstructed tetrode-track showed that this penetration was above the lateral ventricle (Fig. 1A, bottom), in the hippocampal formation (Hp) (Atoji and Wild, 2006).

**Figure 1.**
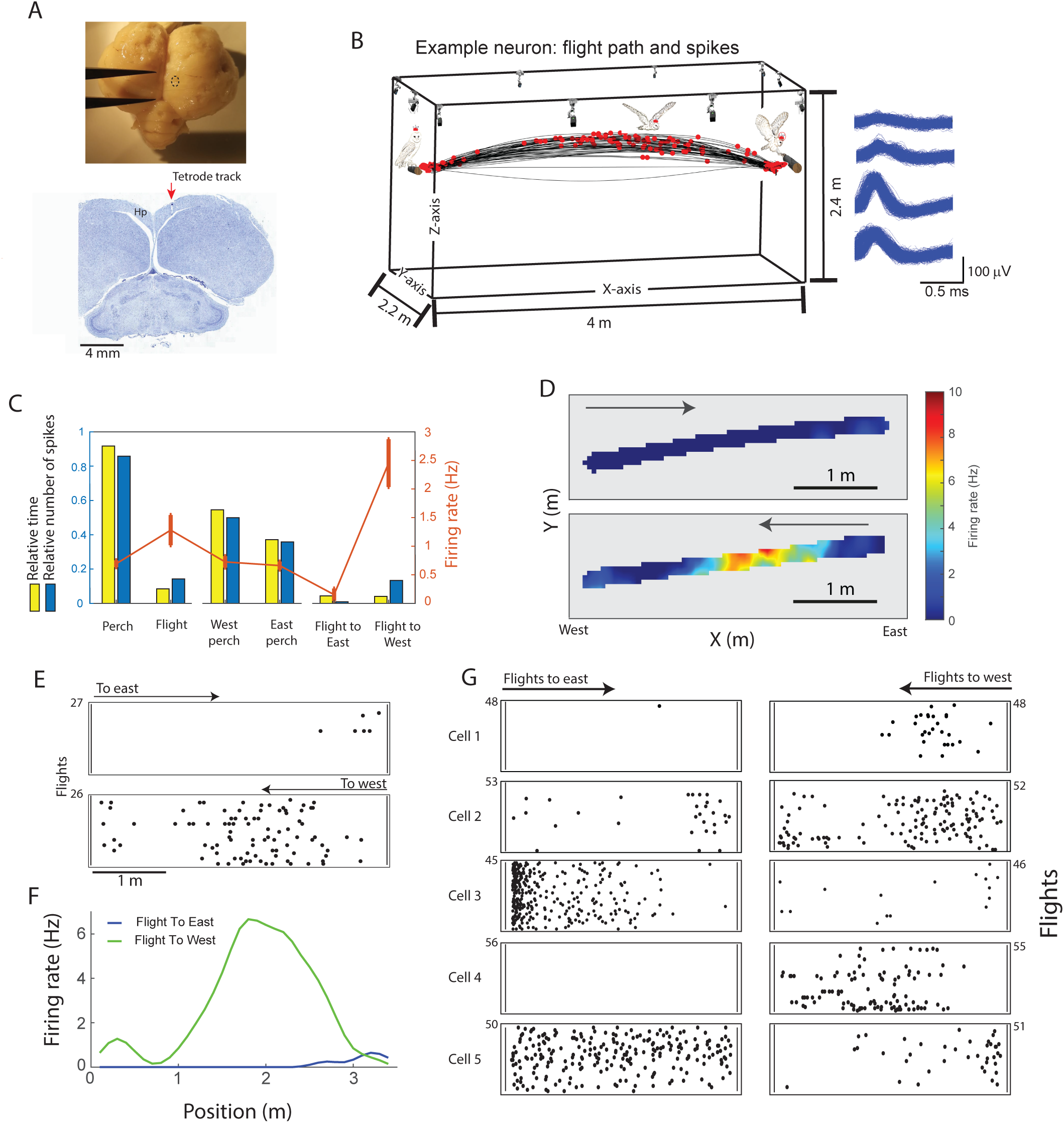
Recording location and example place cells in the hippocampus (Hp). **A.** An upper view of the brain of owl DK following perfusion and removal. The dashed oval marks the position of the craniotomy. The caliper is set to 5 mm. The lower image shows a Nissl stained coronal section of the above brain showing the tetrode track (red arrow). **B-F**. Results from one example neuron. **B.** The flight trajectories of an example recording session (black; shown are both flight directions). Red dots represent the spikes of a single unit whose spike shapes are shown in the inset on the right. The flights are superimposed on an illustration of the flight room and cameras. **C.** The relative time (from the total time of the session) and the relative number of spikes (from the total number of spikes fired by this example neuron) are shown for each of the six possible behavioral states: standing on a perch, flying, standing on the west perch, standing on the east perch, flying eastwards, and flying westwards. The red curve depicts the firing rates in the above six different states (mean ± SE). **D.** A 2D firing-rate map of the cell (top view, XY) drawn separately for westbound flights (bottom) and eastbound flights (top). Arrows mark the flight direction. Colors denote the firing rate. Pixels in which the owl did not visit are shown in gray. **E.** Raster plots showing the spike times along the flight path, for the same cell. The upper raster shows all eastbound flight and the lower raster all westbound flights. Each row is a different flight; the number of flights in each direction is indicated on the y-axis. The x-axis, here and in all examples that follows, is the x-position in the room. **F.** The smoothed firing-rate map of the neuron as a function of position, shown separately for eastbound flights (blue curve) and westbound flights (green curve). **G.** Raster plots for five additional example neurons during flights to the east (left column) and to the west (right column); the number of flights is indicated on the y-axis. Firing-rate maps of all these example neurons were significantly place-tuned and also significantly directionally tuned.

During the ∼20-min recording sessions, the owls flew back and forth between two perches, performing 30–45 flights in each direction, with highly reproducible trajectories that formed a 1D “flyway” (Fig. 1B, Movies 1 and 2) – allowing to statistically compare the firing-rates between flight directions or between perch locations. In the example session shown in Figure 1B-F, the owl spent about 90% of the session standing on either of the perches and about 10% of the time flying between perches (Fig. 1C). Firing rates of the example neuron shown in Figure 1B-F were higher during flight compared to standing (Fig. 1C, red curve). Interestingly, during flight, most of the firing of this neuron occurred in a place-field located between 1.5 to 3 m from the west wall (Fig. 1D) – and moreover, the firing-rate and spatial pattern were clearly direction-dependent, with higher activity during westbound flights compared to eastbound flights (Fig. 1D-F). In this neuron, there were significant differences in firing-rates between perching versus flying and between westbound-flights versus eastbound-flights (bootstrap, p<0.001) – but no firing-rate differences was found between standing on the west perch versus standing on the east perch (p=0.79). Importantly, the neuron exhibited significant place-tuning during westbound flights (p<0.001, compared to spike-shuffling) and the spatial information under uniform coverage assumption (SI_u_; see Methods) was larger than 0.3 – i.e. this was a significant place-cell. Additional examples of significant hippocampal place cells are shown in Figure 1G and Figure S1.

Overall, we identified in Hp three spatially related parameters that significantly modulated the firing rates of the cells: (1*) Place tuning: Selectivity to a specific position in the flight path, in at least one of the directions*. We used the spatial information under uniform coverage assumption, SI_U_, to quantify the place-tuning of the neurons during flight. Out of the 292 cells recorded in Hp, 136 (47%) were significantly place-tuned in at least one direction (rigid circular spike-shuffling, p < 0.01). Among these, 84 neurons (29%) passed our criterion for place cells (SI_u_>0.3): 65 cells were significantly place-tuned during westbound flights and 43 cells during eastbound flights (see examples in Fig. 1E and 1G, and Fig. S1). (2) *Directionality: Flying towards the East versus flying towards the West.* Out of the 292 cells recorded in Hp, 81 cells (28%) exhibited significant directionality (bootstrap, p<0.01; examples in Fig. 1E and 1G, and Fig. S1): 25 cells significantly preferred eastbound flights and 56 cells significantly preferred westbound flights. (3) *Selectivity to standing on east perch versus standing on west perch.* Out of the 292 cells recorded in Hp, 112 cells (38%) exhibited significant selectivity to one of the perches: 52 cells significantly preferred the west perch and 60 cells significantly preferred the east perch (bootstrap, p < 0.01; examples in Fig. 2A). The three types of spatial modulation (place tuning, directionality, and perch selectivity) were intermixed: 28% of the spatially modulated cells displayed more than one type, with some cells’ firing-rate significantly modulated by all three types (n=21; Fig. 2B). Interestingly, among the cells that showed both significant directionality and significant perch sensitivity, there was almost no correlation between the preferred perch and the preferred flight direction (Fig. 2C; Pearson r = 0.26, p = 0.1).

**Figure 2.**
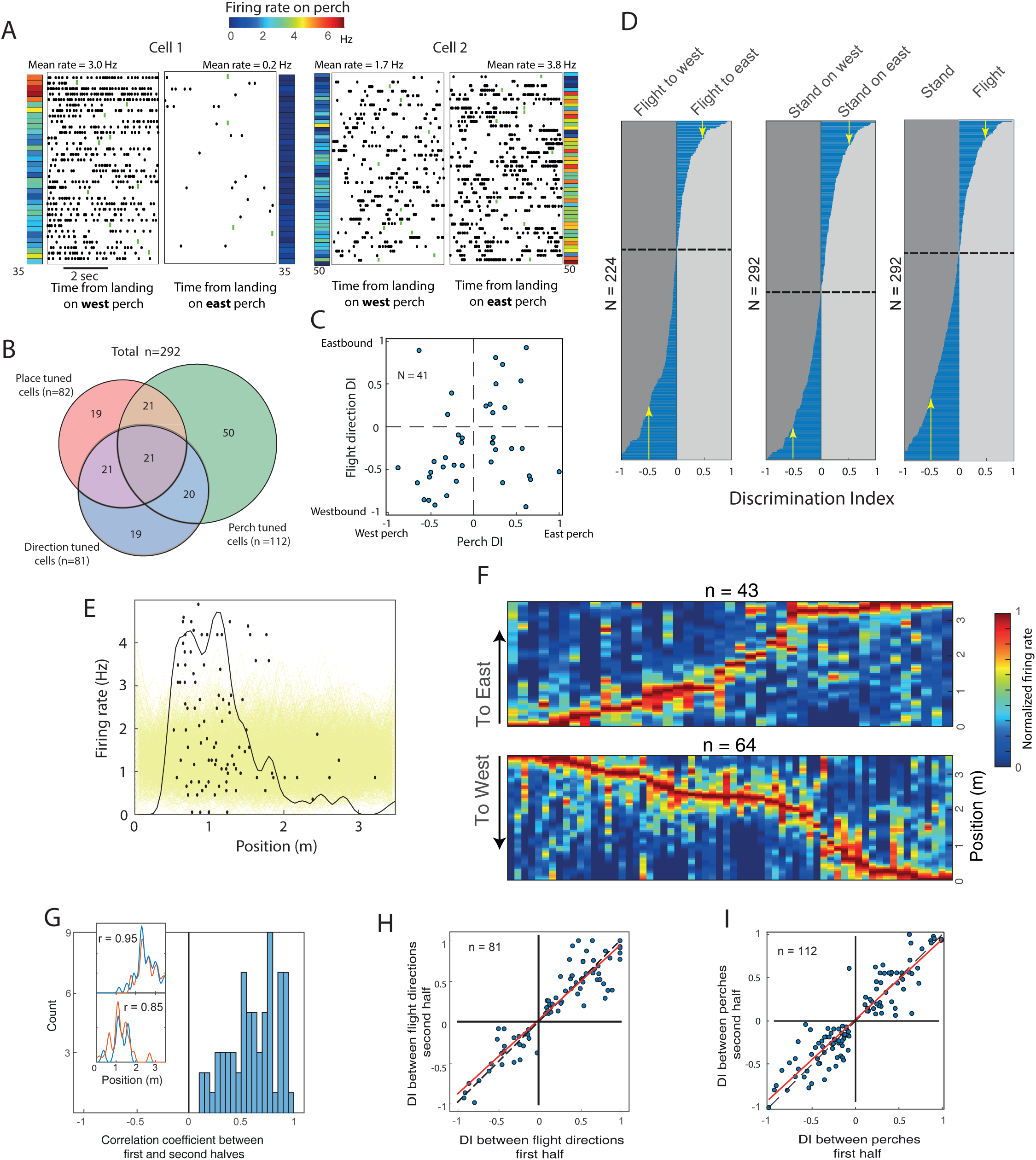
Spatial modulation of neuronal responses in the Hp. **A.** Two example neurons showing significant asymmetric firing between the two perches. Black dots show spike-times relative to the time of landing on the west perch (left raster) or on the east perch (right raster). Shown are the first 5 seconds after landing; in trials where the owl stayed on the perch less than 5 sec, the time of leaving the perch is marked by a green tick. The color-bars on the left and right sides of the rasters designate the mean firing rates for each trial on the west and east perch, respectively. **B.** Venn-diagram showing the relations between three types of spatial modulations (place-tuning during flight, direction selectivity, and perch selectivity) in the population of neurons from the Hp. **C.** A scatterplot showing the relationship, within cells, between the perch discrimination index (DI) and the flight-direction discrimination index. Discrimination index is defined as firing-rate (FR) in one condition minus the other condition, divided by the sum: (FR_1_ – FR_2_) / (FR_1_ + FR_2_). Positive discrimination indices indicate higher firing-rates on the east perch or on eastbound flights, and vice versa. **D.** Histograms showing the discrimination indices for flight direction (left), standing perch position (middle) and standing versus flying (right), for of all cells recorded in the Hp. X-axis, discrimination index; Y-axis, cells: sorted from the most negative DI to the most positive DI. The horizontal dashed line indicates the zero DI. The number of cells in each graph is shown on the left. Flight DIs from cells which fired less than 20 spikes during flight were not included, and hence the number of cells in the left graph is smaller than in the other graphs. Vertical arrows mark cells with DI > 0.5 in both directions. **E.** Example cell showing raster (black dots) and 1D firing-rate map (black curve) for one flight-direction. The yellow curves show 100 firing-rate curves generated from rigidly shuffling the spike trains in-flight in the same flight-direction. **F.** Color plots of the smoothed firing-rate maps (place tuning curves; we used 10-cm spatial bins and Gaussian smoothing with σ = 1.5 bins), for all the place-cells in the Hp (SI_u_ larger than 99% shuffling and SI_u_ > 0.3). Each column depicts a single firing-rate map, normalized by the peak firing-rate of the neuron. Results are shown separately for flights to west (bottom plot) and flights to East (top plot). Curves are sorted from left to right according to the position of the peak firing. The arrows mark the beginning of flight and the flight direction. **G**-**I**. Stability of spatial representations in Hp. **G.** Distribution of Pearson correlation coefficients between the 1D firing-rate maps generated from the first and second halves of the session (firing-rate maps computed separately for each flight-direction). Inset: two example neurons showing the 1D firing-rate maps for the first half of the session (red lines) and second half of the session (blue lines). Pearson correlation coefficient between the red and blue curve is indicated for each example. **H.** The discrimination indices between flight directions, calculated from the first half of the session, are shown versus the discrimination indices calculated from the second half of the session. Only cells that significantly discriminated between flight directions were included in this graph (n = 81 cells). Dashed line, identity-diagonal; red line, linear-regression. **I.** Same as in G but showing the discrimination indices between perches in the first half versus the second half of the session (n = 112 cells).

Across the population of recorded neurons, the normalized differences between the firing-rates during westbound and eastbound flights (flight discrimination indices [DI]; see Methods) ranged from cells strongly preferring westbound flights to cells strongly preferring eastbound flights, with more cells preferring the former (Fig. 2D, left panel; 139 negative DIs compared to 85 positive DIs; Sign test, p<0.001). Likewise, perch preferences varied from west perch preferring cells to east perch preferring cells (Fig. 2D, middle panel). In addition, 54 cells showed significantly higher firing-rates during flight compared to when standing on the perch, and 85 cells showed the opposite (bootstrap, p <0.01; Fig. 2D, right panel).

To examine the spatial distribution of the significant place-fields during flight, we computed the smoothed 1D firing-rate curves for all the significant place-cells (example in Fig. 2E, black curve). At the population level, the place fields covered the entire flyway, with an over-representation of the regions near the perches (Fig. 2F; firing-rate curves were sorted according to the position of the peak firing-rate – separately for the two directions).

To assess the stability of the place-fields, we computed the Pearson correlation coefficient between 1D firing-rate maps that were constructed separately for the first versus second half of the session (Fig. 2G; computed for all flight-directions that exhibited significant place-tuning, n = 191; note that a place-cell can be place-tuned in both directions or only in one direction, hence the number of place-tuned directions is larger than the number of place-tuned cells). The correlations were highly positive (median Pearson correlation coefficient: r = 0.63, n = 69) – indicating stability of the place-tuning. To assess the stability of the neurons’ directional preferences over time, we divided the session into two halves and analyzed the discrimination indices between flight directions and between the two perches. This analysis included only cells which showed a significant flight-direction preference (n = 81) or perch-location preference (n = 112). Discrimination indices of flight direction and perch location were significantly and highly correlated between the two halves of the session (flight direction preference: Pearson r = 0.923, p < 0.0001; perch preference: Pearson r = 0.92, p < 0.0001; see Fig. 2H and 2I). Overall, all spatial preferences in Hp were maintained stably during the experimental session.

### Spatially modulated cells in the HA_p_

In a second set of experiments, we targeted the posterior part of the hyperpallium apicale (HA_p_) – located immediately anterior to the hippocampus. The tetrodes were implanted 5-6 mm anterior from the cerebellum and 2-3 mm lateral from the midline. A reconstructed tetrode-track obtained in one of the owls showed that this penetration was lateral to the edge of the lateral ventricle (Fig. 3A), in an area considered to be in the posterior hyperpallium apicale (HA_p_) (Atoji and Wild, 2006).

**Figure 3.**
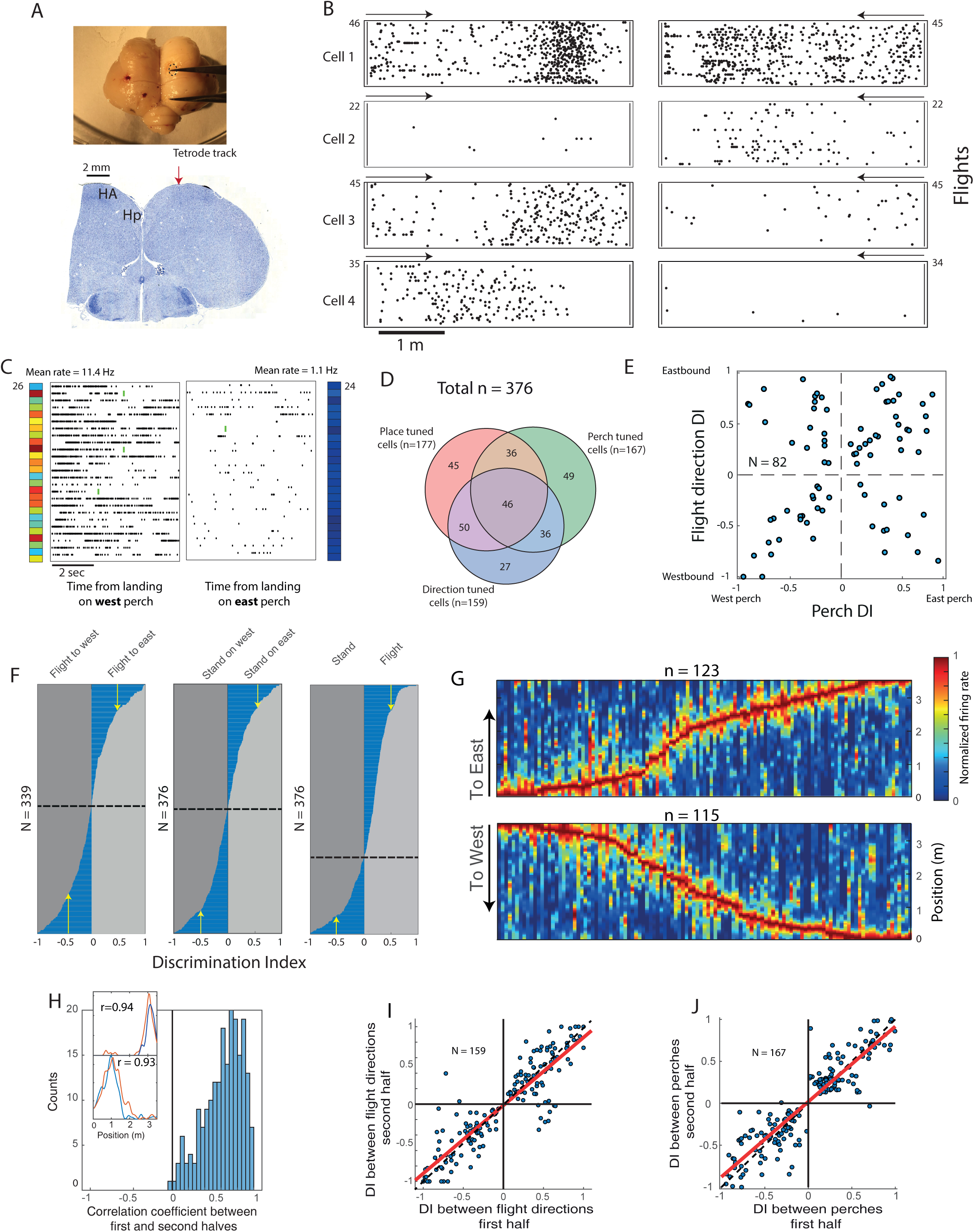
Spatial tuning in the posterior hyperpallium apicale (HA_p_). **A.** The brain of owl TLV1 following perfusion and removal. The dashed oval marks the position of the craniotomy. The caliper is set to 5 mm. The lower image shows a Nissl stained coronal section of the above brain. The tetrode track is marked by the red arrow. **B.** Raster plots of four example cells, showing spike occurrences along the flights eastward (left plots) and flights westward (right plots). Arrows indicate the starting position and direction of flight for each of the plots. The numbers on the y-axis indicate the number of flights. **C.** An example of a perch-selective cell in HA_p_. The rasters show the spike times relative to landing on west perch (left raster) and east perch (right raster). Plotted as in Fig. 2A. **D.** Venn-diagram showing the relations between three types of spatial modulations (place-tuning during flight, direction selectivity, and perch selectivity) in the population of neurons from the HA_p_. **E.** Scatterplot showing the relationship, within cells, between the perch discrimination index and the flight-direction discrimination index. Plotted as in Fig. 2C. **F.** Histograms showing the discrimination indices (DI) of all cells recorded in the HA_p_. Cells were sorted along the y-axis, from the most negative to the most positive DI. Plotted as in Fig. 2D. Vertical arrows mark the cells with DI > 0.5. **G.** Color plots of the smoothed firing rate curves from all place-tuned cells in HA_p_ (SI_u_ larger than 99% of the shuffles and SI_u_ > 0.3). Each column designates a single firing-rate map, separately for flights to West (bottom) and flights to East (top). Curves are sorted from left to right according to the position of the peak firing. The arrows mark the beginning of flight and flight direction. **H**-**J**. Stability of spatial representations in HA_p_. **H.** Discrimination indices between flight directions, for the first half of the session versus the second half of the session. Only cells that significantly discriminated between flight directions were included in this graph. Dashed line, identity-diagonal; red line, linear-regression. **I.** Same as in H but showing the discrimination indices between perches, in the first half versus second half of the session. **J.** Distribution of Pearson correlation coefficients between the 1D firing-rate maps generated from the first half versus second half of the session. Inset: two examples neurons, showing the 1D firing-rate maps for the first half of the session (red lines) and second half of the session (blue lines). Pearson correlation coefficient between the red and blue curve are indicated for each example.

In the HA_p_ we found the same three types of spatially modulated neurons that we found in the Hp. Surprisingly, however, a larger percentage of spatially tuned neurons were encountered in the HA_p_ as compared to the Hp: (1) *Place tuning:* Out of 376 cells, the firing rates of 280 cells (74%) were significantly modulated by the position along the flight trajectory in at least one direction (spike-shuffling statistics, 99% percentile: i.e., p < 0.01). Among these, 177 cells (47%) passed the criterion for place-cells (SI_u_ > 0.3; examples in Fig. 3B, and Fig. S2): 116 cells were significantly spatially-tuned during westbound flights and 123 during eastbound flights. (2) *Directionality*: Out of 376 cells, 159 cells (42%) exhibited significant directionality: 74 cells significantly preferred eastbound flights and 85 significantly preferred westbound flights (bootstrap statistics, p<0.01; examples in Fig. 3B, and Fig. S2). (3) *Perch selectivity:* When standing on the perch, 167 out of 376 cells (44.4%) exhibited significant selectivity to one of the perches (example in Fig. 3C): 82 cells significantly preferred the east perch and 85 cells the west perch (based on bootstrap criterion, p < 0.01).

In HA_p_, a substantial overlap between the three types of spatial modulation was observed, with 46 cells exhibiting significant modulation by all three parameters (place-tuning, directionality, and perch-preference; Fig 3D). Again, there was no significant correlation between the preferred perch side and the preferred flight direction (Fig. 3E; Pearson r = 0.17, p = 0.1). In both brain regions (Hp and HA_p_) a substantial number of cells distinguished between flight direction and/or perch position, with discrimination indices larger than 0.5 (arrows in Figs. 2D and 3F). In the HA_p_, no significant bias for flight direction was observed (Fig. 3F, left panel; 165 positive DIs versus 174 negative DIs; Sign test, p = 0.74). Finally, in both Hp and HA_p_ the place-fields exhibited over-representation of space at the two ends of the flyway, near the perches (Fig. 2F and Fig. 3G) – as found in rodents and bats near reward zones (Eliav et al., 2021; Geva-Sagiv et al., 2016; Hollup et al., 2001). As in Hp, also in HA_p_ the neural representations of location (place-tuning), direction, and perch-location were highly stable between the first and second halves of the session (Fig. 3H-J; place-tuning: median Pearson correlation between first versus second half, r = 0.678, n = 205; directionality: Pearson r = 0.88, n = 159; perch preference: r = 0.91, n = 167).

Occasionally, the owls made a U-turn in mid-flight, turning back to land on the perch from which they took off. These events provided an opportunity to examine whether the directional sensitivity is determined by the flight direction *per se*, independent of the starting position. In the example session shown in Figure 4A, the owl made two U-turns. Action potentials in the U-turn trials were more abundant when the owl was flying westbound, along the preferred direction of the cell (Fig. 4A, top panel). In most of the recordings made during U-turn trials, the number of spikes that were discharged when the owl was flying in the preferred direction was larger compared to flying in the opposite direction (Fig. 4B; Sign test, p < 0.0001; n = 584 [pooled data from Hp and HA_p_]). Thus, the neuronal directionality is determined by the instantaneous flight-direction of the owl and not by the takeoff direction.

**Figure 4.**
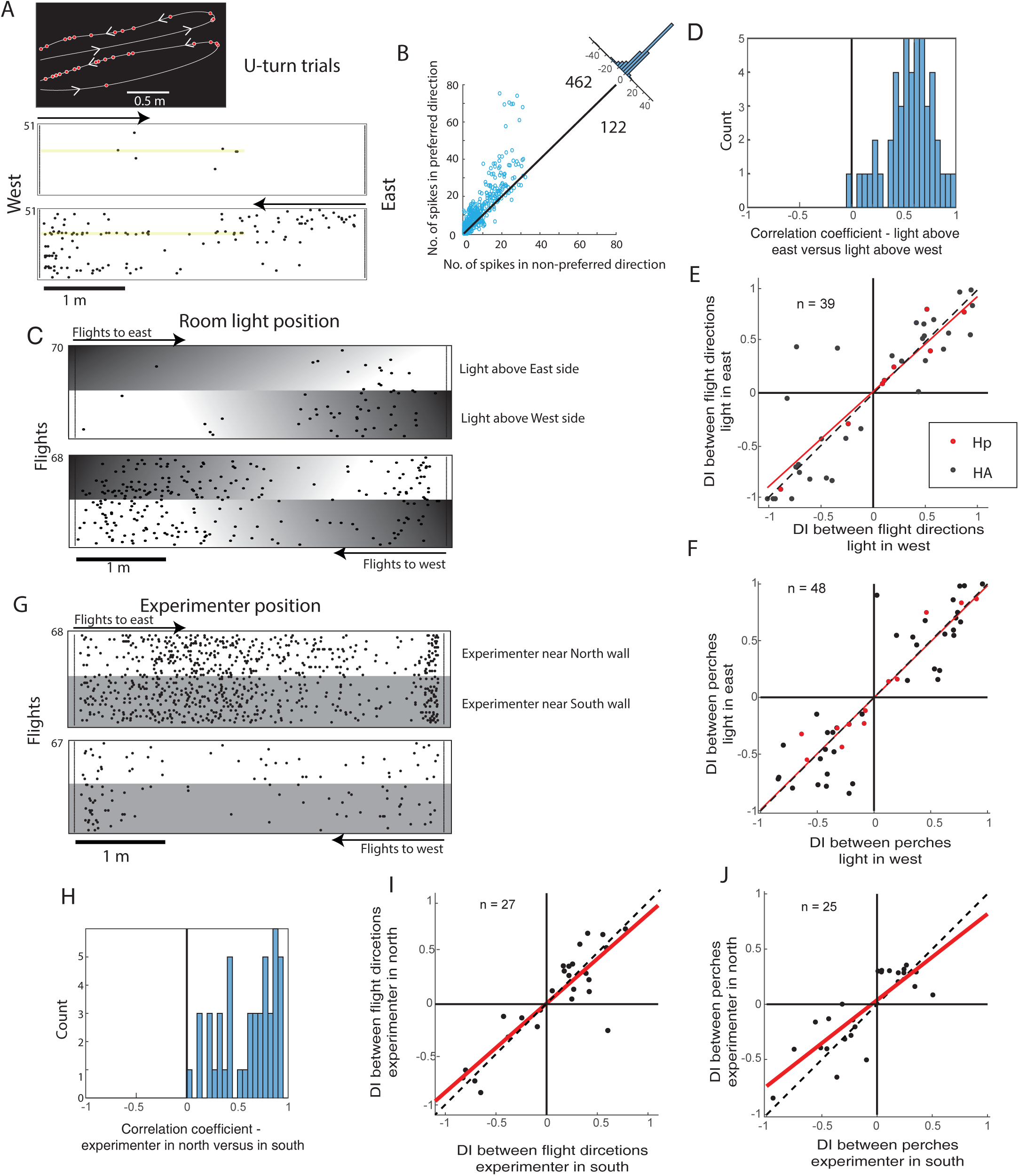
U-turns and experimental manipulations. **A**-**B**. U-turns. **A.** Raster-plots for an example neuron recorded in HA_p_, showing significant preference to westbound flights (bottom) over eastbound flights (top). The yellow lines on the rasters mark the two trials in which the owl started on the west perch and made a U-turn in mid-flight and landed again on the west perch. Inset above (black): top view of the flight trajectories of these two trials. The trajectories were shifted on the Y-axis to separate them for display purposes. Red dots on the trajectories indicate spikes. The arrowheads point to the direction of flight. **B.** Scatter-plot of the number of spikes in the preferred direction versus the non-preferred direction of the neurons, for all the U-turn trials. The number of dots above and below the diagonal are indicated. Data are pooled together from Hp and HA_p_ . **C**-**J**. Experimental manipulations. **C.** An example of a neuron’s firing during flights when the light was above the west perch and then flipped in the middle of the session to the opposite side of the room. The direction of the lighting pattern in the room is indicated by the white-to-black background shading. Bottom raster shows flights to west and top raster flights to east. **D.** Distribution of Pearson correlation coefficients between the 1D firing-rate maps computed for the two parts of the session: light-in-west versus light-in-east. **E.** Scatter-plot showing the discrimination indices (DI) between flight directions when the light was in the east versus the discrimination indices when the light was in the west. Red dots, cells that were recorded in Hp; blue dots, cells recorded in HA_p_. Dashed line, identity-diagonal; red line, linear-regression. **F.** Same as E but showing the discrimination indices between the perches when the light was in the east versus when the light was in the west. **G**. Raster plots of an example neuron recorded in a session during which the experimenter switched sides in the middle of the session: the experimenter stood first adjacent to the south wall and later adjacent to the north wall. Plotted as in C. Gray areas in the raster mark the trials in which the experimenter was in the south, and white areas – in the north**. H-J.** Graphs plotted as D-F, but showing the Pearson correlations between 1D firing-rate maps when the experimenter was in the south versus north (H), the discrimination indices between flight-directions when the experimenter was in the south versus north (I), and the discrimination indices between perches when the experimenter was in the south versus north (J). In the experiment in H-J, all the neurons (dots) were recorded in HA_p_.

Next, we examined the effect of lighting conditions on the place-tuning of neurons. The experimental room was normally illuminated by three LED lamps: one at the center, one above the east perch and one above the west perch. In several of the sessions we changed the illumination in mid-session: in the first half of the session the arena was illuminated only with the light above the west perch, and in the second half we switched to illuminating only with the eastern light. This created an asymmetric lighting in the room that was switched to the opposite side in mid-session. The switch in lighting-conditions had little effect on the firing pattern of the example cell shown in Figure 4C – and importantly, it changed neither the place-tuning nor the flight-direction preference. At the population level, the pattern of place-tuning (firing-rate curves) was stable independent of the lighting condition for most of the significant place-cells tested with this manipulation (Fig. 4D; median Pearson correlation coefficient: r = 0.58, n = 47 tuning curves from 29 cells). Flight direction preference was not changed in most of the direction-significant neurons which were tested in this experiment (Fig. 4E; Pearson r = 0.87, p < 0.0001, n = 39), and the perch preference was also not changed (Fig. 4F; Pearson r = 0.86, p < 0.0001, n = 48).

During the recording session, an experimenter was present in the room with the owl, normally adjacent to the South wall (see Methods), thus serving effectively as a stable “landmark” – which was highly salient for the owl. To test the influence of the experimenter’s position, in several of the sessions the experimenter switched position in mid-session to stand adjacent to the North wall. This switch, had no obvious effect on the firing properties of the example cell shown in Figure 4G. At the population level, we found that – similar to the switch in lighting conditions – place-tuning was maintained independent of the experimenter’s position (Fig. 4H; median Pearson correlation coefficient: r = 0.72, n = 47), as was the flight-direction preference (Fig. 4I; Pearson r = 0.64, p < 0.0001, n = 27) and perch preference (Fig. 4J; r = 0.62, p < 0.0001, n = 25).

Taking together the experimental switching of the light-source position and experimenter’s position, these results suggest that – as in rodent place-cells (O’Keefe and Nadel, 1978) – owl place-cells are not sensitive to any one particular individual landmark (we note that many visual landmarks were present in the room: see Movie 1). Instead, place-cells encode the animal’s position in a more abstract manner, likely based on a combination of numerous landmarks.

### Single unit responses in visual Wulst

To examine whether the newly found spatial representation in the HA_p_ is restricted to the posterior part of the HA – adjacent to the Hp – or is anatomically widespread across the avian hyperpallium, we targeted our recordings in the next experiment to the central part of the hyperpallium: the visual Wulst. Visual Wulst in barn-owls is a well-studied, retinotopically organized visual area that is the main recipient of the thalamo-fugal visual pathway (Nieder and Wagner, 2000; Pettigrew, 1979). In this location, some of the neurons displayed, in a head-fixed condition, small visual receptive fields (RF, ∼5° diameter) at the frontal visual field (Fig. 5A). Based on previous mapping of visual Wulst in barn owls, such RFs correspond to a location at the lateral central part of the retinotopic visual map in Wulst (Pettigrew and Konishi, 1976).

**Figure 5.**
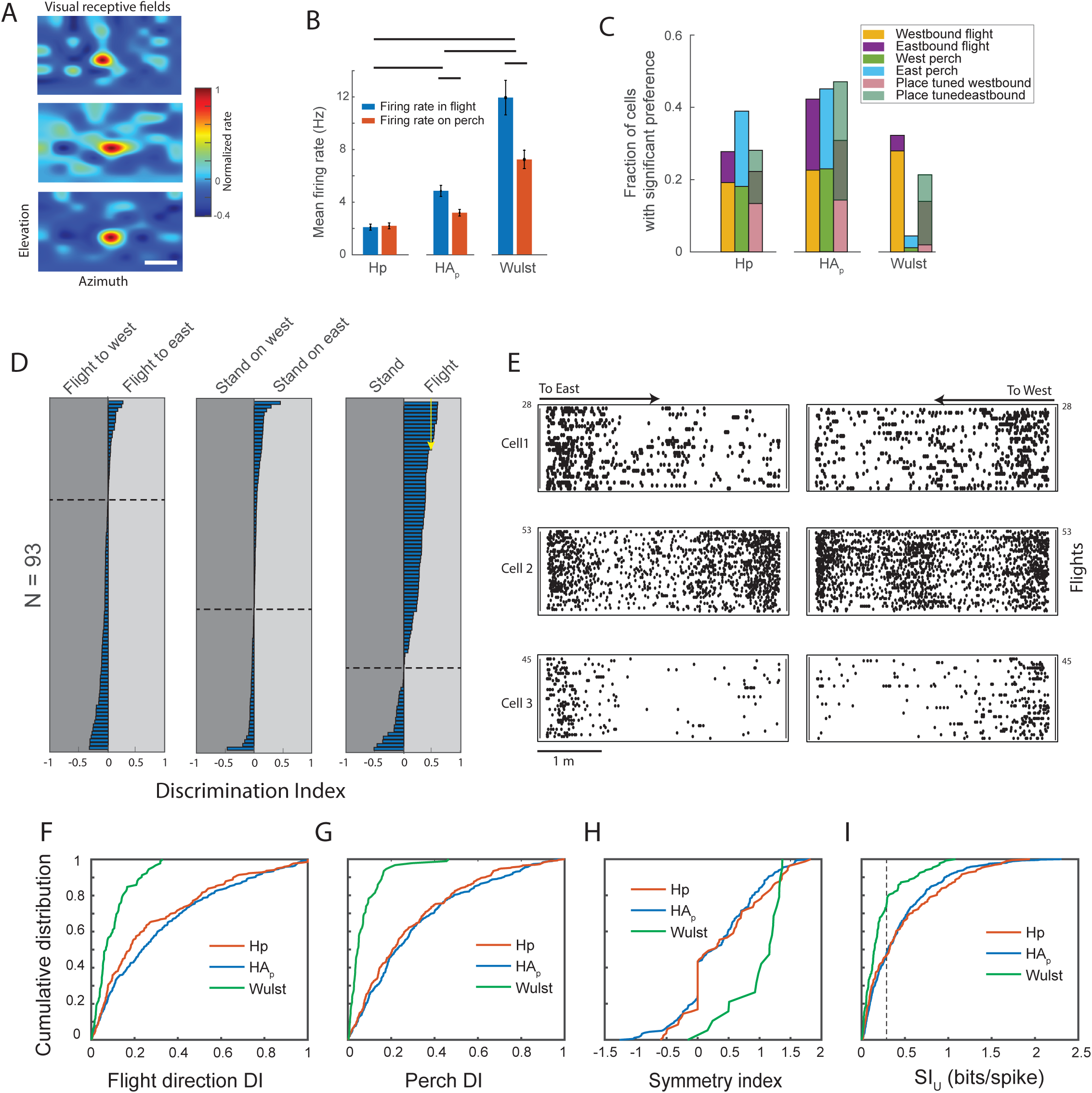
Recordings from visual Wulst, and comparison between Hp, HA_p_ and Wulst. **A**. Visual receptive fields of three neurons recorded in visual Wulst. Color plots show the smoothed normalized responses (firing-rates after stimulus onset minus firing-rates before stimulus onset) as a function of the position of the visual stimulus on the screen. Scale bar, 10°. **B.** Comparison between the mean firing-rates of all the neurons recorded in Hp, HA_p_ and Wulst, separately for firing-rates during flight (blue) and firing rates when standing on the perch (red). Error bars are SEM. Horizontal lines above the bars indicate p < 0.01 (*t*-test). **C.** Comparison between the fractions of cells in Hp, HA_p_ and Wulst that significantly (p < 0.01) preferred flights to east or flights to west, and east perch or west perch. The third bar in each brain-area shows the fraction of place-cells (neurons with SI_u_ that was significant compared to spike-shuffling with p < 0.01 [99% shuffling] and SI_u_ >0.3), during westbound flights or eastbound flights. The middle region in each bar indicates the cells whose firing rate was place-tuned in both directions. **D.** Summary of the discrimination indices between flight-directions (left plot), perch locations (middle plot) and flight versus standing (right plot), for all the neurons recorded in visual Wulst (n = 93). Plotted as in Fig. 2D. The y-axis represents the neurons, sorted by the discrimination-index. Vertical arrow in the right plot marks the neurons with values large than 0.5; notably, in the left and middle plots all values were smaller than 0.5. **E.** Examples of spike-rasters during flights for three cells in Wulst whose firing-rates were significantly modulated along the flight. Left column, spikes during eastbound flights; right column, spikes during westbound flights. Arrows indicate the beginning and direction of flights for each raster. Numbers on the y-axis designate the number of trials in each raster. **F.** Cumulative distributions of the discrimination indices (DI) between flight directions. Results are from the population of neurons that significantly discriminated between directions in the three brain regions: Hp, HA_p_ and Wulst. **G.** Same as in F but between perches. **H.** Cumulative distributions of the symmetry indices from all place-tuned neurons in the three brain regions. **I.** Cumulative distributions of the spatial information indices (SI_U_) from all spatially modulated firing-rate maps (spike-shuffling statistics; p <0.01), in Hp, HA_p_ and Wulst. The dashed-line marks the value used as a criterion for place-tuned cells (SI_u_=0.3).

We then turned to recording visual Wulst neurons in flight. In total, we recorded 93 single units in Wulst from two owls (see Table S1). The average firing-rate during flight was significantly higher than during perching (Fig. 5B; *t*-test, p=0.0076). Moreover, Firing-rates in Wulst were significantly higher compared to firing-rates in both Hp and HA_p_ (Fig. 5B; *t*-test, p<0.001), and firing-rates in HA_p_ were significantly higher than in Hp (Fig. 5B; *t*-test, p<0.001). In Wulst, as in Hp and HA_p_, we found neurons that significantly discriminated between the two flight directions (Fig. 5C; 32% of the neurons discriminated between flight-directions; examples in Fig. 5E and Fig. S3). However, unlike in Hp and HA_p_, discrimination between perches was very weak in visual Wulst (only 4 cells significantly discriminated between perches). Moreover, neurons in Wulst showed strikingly lower discrimination indices for flight-direction and perch-position, as compared to HP/HA_p_: While in HP and HA_p_ a substantial number of neurons had high discrimination indices (discrimination indices higher than 0.5 are marked by arrows in Figs. 2D and 3F), the maximal discrimination index in Wulst was 0.33 for flight-direction and 0.38 for perch-position (Fig. 5D). The DIs, both for flight direction and perch position, were significantly smaller in Wulst as compared to Hp and HA_p_ (Figs. 5F and 5G; Kolmogorov-Smirnov test, p <0.001). In other words, cells that showed substantial differences between firing-rates in different directions or perches were found in Hp and HA_p_ – but not in Wulst. Another difference was that selectivity in Wulst tended to be non-uniform: Among the cells that significantly discriminated between directions, 26 preferred westbound flights and 4 eastbound flights (Table S1, Fig. 5C)

In Wulst, 20 cells out of 93 (22%) passed the criterion for place cells (spike-shuffling statistics, p < 0.01 and SI_u_ > 0.3). This compares with 84 place cells out of 292 in Hp (29%) and 181 place cells out of 376 in HA_p_ (47%) (Fig. 5C). However, unlike in Hp and HA_p_ where the firing patterns of the place-cells were mostly direction-dependent, the 20 place-cells in visual Wulst were highly symmetrical between the two flight directions – i.e., the firing pattern in one direction was a mirror image of the firing pattern in the opposite direction (Fig. S4 and examples in Fig. 5E and Fig. S3). The cumulative distribution of the symmetry indices was significantly positive in Wulst compared to Hp and HA_p_ (Kolmogorov-Smirnov test, p <0.001; Fig. 5H) – indicating that many Wulst neurons encoded the distance flown from takeoff, regardless of flight direction.

Finally, the values of the spatial information, SI_U_, were significantly smaller in Wulst compared to HA_p_ and Hp (Kolmogorov-Smirnov test, p < 0.001; Fig. 5I; using all neurons with significant spike-shuffling statistics of p<0.01 on place-tuning, but without applying the criterion of SI_u_ > 0.3). The reduced magnitude of place-tuning in Wulst, as compared to Hp and HA_p_, was significant when using three common indices to quantify spatial tuning in hippocampal neurons (Skaggs et al., 1996): spatial information, sparsity, and selectivity (Fig. S6) – suggesting that, as in mammals, strong allocentric place-tuning is found in neurons of the owl’s hippocampal formation and adjacent regions, but not in primary visual areas.

## Discussion

Single unit recordings in freely behaving animals revolutionized the study of the neural mechanisms of behavior (O’Keefe and Nadel, 1978; Ziv and Ghosh, 2015). Technological advances in the last two decades allowed recording single units from flying animals, including recordings in the hippocampus of flying bats (Yartsev and Ulanovsky, 2013). Here we report the results of single-unit recordings in the hippocampus of flying barn owls. To our knowledge this is the first report of single unit recording during flight in the hippocampus of any species of bird.

Barn owls are nocturnal central-place predators that spend much of the night-time perching on strategic posts, occasionally leaving the branch to explore for rodents (Roulin, 2020; Rozman et al., 2021). In an enclosed space, they tend to fly from one standing position to another in straight lines and do not cover the space like bats or some other species of birds do. Therefore, in our experiments, we explored neuronal firing during a back-and-forth flight between two perches on opposite walls. This 1D behavior is comparable to a linear track behavior (Las and Ulanovsky, 2014). Multiple studies investigated the firing of place-cells in animals walking back-and-forth along a linear track, and have reported spatial tuning that differs between both running-directions (e.g., McNaughton et al., 1983; O’Keefe and Recce, 1993; Ziv et al., 2013). Directionality of place tuning was found also during back-and-forth 1D flying in bats (Eliav et al., 2021; Geva-Sagiv et al., 2016). Similarly, place cells recorded recently in the titmouse (a food-caching bird) showed directional place-fields in a linear track behavior (Payne et al., 2021). Many of the neurons recorded in the barn owl showed a similar behavior: namely, their firing-rates were significantly modulated by the position along the path, and this place-tuning was different for each direction. Spatial and directional modulations were stable during the recording session and were robust to salient changes in the lighting and in the position of the experimenter in the room. Therefore, our findings suggest the existence of place-cells in the dorsal pallium of barn owls – both in the hippocampal formation (Hp) and in the adjacent part of the hyperpallium apicale (HA_p_). In addition, some of the cells robustly discriminated between the perches while the owls were standing in-between flights – perhaps similar to hippocampal CA2 neurons in rats that were reported to encode the rat’s position during immobility (Kay et al., 2016).

Although our findings are consistent with place-cells, there are other possibilities that should be considered. One possibility is that we are recording from visual neurons that are sensitive to a specific visual cue that passes through a visual receptive-field. Since the flights were stereotypic, a certain cue in the room is expected to appear consistently at a certain location and direction (Carandini et al., 1998). However, our findings do not support this possibility. First, different cells recorded in the same owl showed opposite direction sensitivities (Table S1). Visual RFs in the hyperpallium of barn owls are organized retinotopically to map the contralateral side (Pettigrew, 1979). Therefore, if these were classical visual neurons, the responses of neurons in a single side of the brain are expected to be spatially correlated, i.e. display similar directional tunings – which was not the case. Second, switching lighting direction in the room did not correspondingly switch the directional preference. Third, when the most salient visual cue in the room – the experimenter – switched sides, the directional preference was maintained. Fourth and finally, we manually projected an ophthalmoscope light on the owls’ eyes and searched for visually evoked responses: clear visual evoked responses were apparent in the recordings from Wulst, but not in the other recordings. Recordings from Wulst, a well-known primary visual area, showed cells that were significantly modulated by position and direction – but less than in Hp and HA_p_. Moreover, place-tuned neurons in Wulst showed highly symmetrical responses between the two flight directions (Fig. S3 and S4). Symmetrical responses, in rodents, have been related to distance-coding and not place coding (Ravassard et al., 2013). The symmetrical spatial modulation in Wulst may reflect sensitivity of the visual neurons to the changes in optic flow during flight, which are expected to be correlated with distances from the incoming walls (Serres and Ruffier, 2017).

Another possibility is that the place-cells in Hp and HA_p_ are time-cells – namely, they are modulated by the time from flight onset rather than the location. Time cells, which fire at specific times from the initiation of a behavior, have been found in the hippocampus of rodents (Eichenbaum, 2014). In our experiments, the flights were stereotypic and all followed similar velocities (Fig. S6). Therefore, it is difficult to dissociate between temporally and spatially modulated cells during flight. However, firing during U-turn trials (Fig. 4A and B) were consistent with place coding rather than coding the time from take-off. Moreover, analysis of spike rates during standing on the perch showed that the firing rates of the majority of cells were stable during the first 10 seconds following landing on the perch (Fig. S5). Only a few cells fired significantly stronger at particular time-moments after landing – which could be evidence for a small subpopulation of time-cells in the bird hippocampal formation (see in particular the cell in Fig. S5E).

In mammals, place cells are found predominantly in the hippocampus. Here, we found robust spatial modulation of neurons both in the owl hippocampus (Hp), and in an adjacent region – the posterior part of the hyperpallium apicale (HA_p_). The HA, including the HA_p_, is the upper layer of the hyperpallium. In one leading theory, the HA is comparable to neocortical layers V-VI (Stacho et al., 2020). The anterior part of the hyperpallium is a somatosensory area (Wild, 1997) and posterior to it is a primary visual area (Wulst) (Karten et al., 1973b; Pettigrew, 1979). However, the function of the most posterior and medial areas of the hyperpallium is unknown (Karten et al., 1973a). Moreover, the functional borders and subdivisions of Hp and HA are not clearly defined (Herold et al., 2019). It is therefore difficult to compare the anatomy of our findings in HA_p_ with that of mammals. One possibility is that the most posterior parts of the hyperpallium are functionally related to the neighboring Hp and together comprise a network of spatial representation. Interesting in this respect is the finding in pigeons of a belt in HA_p_ that is densely stained with substance-P positive cells (Erichsen et al., 1991). This distinct nucleus (named Spf) was suggested as a possible candidate for an entorhinal cortex analogue in birds (Székely, 1999). It is noteworthy also that although in mammals place cells have been primarily described as a hippocampal phenomenon, spatial modulation of firing-rates has been found also in several neocortical areas (Diamanti et al., 2021; Esteves et al., 2021; Long and Zhang, 2021). Thus, our finding of place-cells in HA_p_ may be in-line with findings from mammals.

The study of hippocampal place cells in mammals has focused primarily on rats and mice, and more recently also bats. The anatomical and physiological similarities of place cells, grid cells and head-direction cells among these mammalian species are striking (Las and Ulanovsky, 2014; Geva-Sagiv et al., 2015). However, because of the limited number of species studied to date, it is not clear to what extent the spatial characteristics of hippocampal-formation neurons are correlated with the specific ecological demands of the species – or perhaps they are a general characteristic of mammalian hippocampus. The study of neural representations in the avian Hp is at its infancy relative to mammals. Yet, the accumulating data, together with the results reported here, already covers five distinct bird species, and the emerging view is of substantial variability between species (Ben-Yishay et al., 2021; Kahn et al., 2008; Payne et al., 2021). Titmouses with their unique capability to memorize the locations of thousands of seeds have an enlarged hippocampus as well as an exceptional abundance of place-cells (Payne et al., 2021). Similarly here, in the barn owl – a central place forager that strongly relies on memory to navigate to strategic standing posts and to its roost at night (Rozman et al., 2021) – we found robust place-cell representation. This suggests that, across species, the occurrence of place-cells is correlated with strong reliance on spatial memory. We speculate that the owl’s ability to hunt and navigate in nearly-complete darkness is made possible, in part, by an exceptional hippocampal-based spatial memory – and thus clear place-cells are observed. We believe that it is crucial to further collect additional data from hippocampal neurons across multiple species – in order to elucidate the evolutionary origins and functional properties of place cells.

## METHODS

### Animals

Six adult Barn Owls (*Tyto alba*) were used for the study (four females and two males; Arya, DB, TLV, DB2, DK and BB; see Table S1). The owls were hatched in our in-house breeding colony and were hand-raised from 10 to 60 days of age, and thus were well acclimatized to human presence and handling. All procedures were approved by the Technion’s Institutional Animal Care and Use Committee and were in accordance with the Israeli law for the prevention of cruelty to animals.

### Behavioral setup

Experiments were conducted in a 4 × 2.2 × 2.4 m windowless room, dimly lit by three non-flickering LED light sources. The short walls of the room faced East and West, and the long walls faced South and North. The room contained two wooden standing-perches (1.7 m above the ground, 50 cm long, 10 cm wide, and parallel to the wall), one on the East wall and one on the opposing West wall. At the beginning of each recording session, the owl was released in the room and typically flew to one of the perches to stand there. To encourage flights between the perches the experimenter entered the room and stood adjacent to the south wall (in a few control experiments the experimenter moved in mid-session to the opposite wall). When the owl was standing on one of the perches, the experimenter made a gesture towards the owl, by either stepping slowly towards it or slowly moving the arm on the side of the owl – which encouraged the owl to start flying. This procedure was repeated every day during a recording session of ∼20 minutes. The owls quickly adjusted and learned to respond to small experimenter’s gestures, by flying to the opposite perch, often flying spontaneously back and forth multiple times without any movements by the experimenter. During a session, owls flew on average 82 ± 20 flights (mean ± SD).

### Surgery and neural recordings

Owls were prepared for chronic electrophysiological recordings with a single surgical procedure: the birds were anaesthetized with 2% isoflurane in a 4:5 mixture of nitrous oxide and oxygen. Owls were then positioned in a stereotaxic frame (Kopf, small animals instrument Model 963) using custom-made ear bars. Lidocaine (Lidocaine HCl 2% and Epinephrine) was injected locally at the incision site. The skull was exposed and cleaned. Four skull screws were inserted at the posterior part of the skull and one ground screw was inserted at the right frontal part of the skull. Distances on the skull were measured from the anterior edge of the dorsal neck muscle (tendon attachment position on the skull) – which is the standard cranial landmark used in barn owls – using a fine surgical caliper, to determine craniotomy positions. Coordinates for the hippocampus (Hp): 13 mm anterior and 1.5–2 mm lateral from the muscle. For the posterior part of hyperpallium apicale (HA_p_): 14–15 mm anterior and 2–3 mm lateral from the muscle. For Wulst: 18 mm anterior and 6 mm lateral. A 2-mm diameter craniotomy was drilled around the desired position and a small nick in the dura was made at the center of the craniotomy using a surgical needle. We used a custom-made Microdrive (modified from (Weiss et al., 2017)), containing four tetrodes (made of 17.8 µm platinum-iridium wire, California Fine Wire); the tetrodes were platinum-black plated to reduce the impedance to 500–800 kOhm at 1 kHz. The microdrive was carefully lowered until the tetrodes smoothly entered the brain tissue. Penetration was stopped 400 µm below the brain surface. Antibiotic ointment (Chloramfenicol 5%) was applied to the brain surface, followed by a thin coat of a silicon elastomer (Kwik-Sil). The microdrive was then connected to the ground screw with a silver wire and attached to the skull with adhesive cement (C&B Metabond) and light-cured dental resin (Spident Inc. EsFlow). Meloxicam (0.5 mg/kg) was injected intramuscularly, and the owls were then positioned in a heated chamber to recover.

Neuronal signals were recorded using a 16-channel wireless neural-recording device (‘neural-logger’, Spikelog-16, Deuteron Technologies) which was attached to the Omnetics connector on the microdrive. Signals from all 16 channels of the 4 tetrodes were amplified (×200), bandpass filtered (300 – 7,000 Hz) and sampled continuously at 29.3 kHz per channel. Data were stored on-board the neural-logger. Recording sessions were performed almost every day, typically for a few weeks, until signal-to-noise ratio dropped. In total we conducted 125 recording-sessions from five owls (see Table S1 for details). Tetrodes were advanced by 50 µm every day. If no spikes were detected, tetrodes were continuously lowered until spikes were observed. After reaching a depth of 1,500 µm below the brain surface, in some of the experiments the tetrodes were retracted to regions of previously-observed spiking activity, and additional experiments were conducted. Restricting the recordings to below 1.5 mm from the brain surface ensured that the bulk of the recording was within the Hp layer (above the lateral ventrical) or in the superficial layers of the hyperpallium (hyperpallium apicale [HA]). At the end of the experiment the microdrive was carefully removed, the craniotomy was sealed using Kwik-wil, and a new Microdrive was installed in the opposite hemisphere for further recordings.

### Behavioral tracking

The position of the owl in the room was continuously tracked at a rate of 120 Hz, via eight high-speed infrared cameras (OptiTrack, Oregon, USA) that were placed around the room, which captured the positions of infrared reflectors rigidly attached to the microdrive on the head of the owl. Online 3D reconstruction of the 3D position of the reflectors (accuracy of ∼5mm) was achieved with the Motive software (OptiTrack, Oregon, USA). The room coordinates and the 3D position of the two perches were also recorded, alongside with the owl’s position, by placing infrared markers on the edges of the perches, as well as markers on the floor corner to mark the zero coordinate.

For synchronization between behavioral tracking and neural recordings, signals were sent from the neural-logger control box at a rate of 0.5 Hz with a random jitter of ± 50 ms. Signals were transmitted wirelessly to the neurologger and in parallel to the computer running the Motive program, timestamped and saved. The timestamps in the neural-logger and in the Motive files were synchronized by utilizing polynomial curve fitting between the pulses recorded on both systems.

### Data processing and spike sorting

In Subsequent processing (off-line), electrical recordings were band-pass filtered between 600– 6,000 Hz. The first and last minutes of recordings were discarded to avoid filtering edge effects.

Large amplitude artifacts were detected and removed from all channels based on absolute voltage larger than 0.8 mV. Then an adaptive voltage threshold (thr) was used for spike detection:

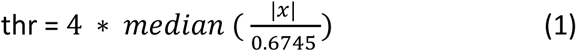

Where x is the filtered recorded signal over a running window of 1 minute, the division by 0.6745 accounts for the relationship between the median of the noise and the standard deviation of a normal distribution. Whenever one channel in a tetrode crossed the threshold value, a 1-ms segment was saved from all channels on the tetrode. Manual spike sorting was performed using the SpikeSort3D software (Neuralynx) and consisted of plotting the spikes in 3D parameter space, finding the features which gave the best cluster separation – primarily spike amplitudes – and manually selecting well-isolated clusters. Coincidence-detection algorithm was used to further identify and remove noise, where equal-amplitude events occurring simultaneously between different tetrodes were removed from analysis. Additionally, all detected spikes were compared to a pre-existing spike-shape database (template matching) by correlating different segments of the spike shape to each template from the database: A spike was included for further analysis if its Pearson-correlation coefficient with any of the spike templates was larger than 0.8. Spikes were categorized as multi-unit or single-unit, taking into account their clusters, spike shapes, and a refractory period (<2 ms) in the interspike-interval histogram. Spikes considered multi-units were not analyzed. Single units with less than 50 spikes in a session were excluded.

### Data analysis

Analyses of all the behavioral and neural data were done using custom Matlab code. To classify the behavior to perching versus flying, the 3D head trajectory was smoothed (spline smoothing) and the tangential velocity was calculated. A tangential velocity higher than 80 cm/s was classified as flight. In addition, short epochs of velocity lower than 80 cm/s during flight (in U-turns or during short hovering before landing) were included as flights if they happened further than 10 cm from the perch. Epochs of velocity higher than 80 cm/s recorded at a distance of less than 10 cm from the perch were discarded from the analysis. These criteria imply that flight take-off and landing were not analyzed. Flight direction was classified to westbound or eastbound, based on the *x*-direction of the flight velocity (where *x* is the 4-meter long axis of the room, which connected the two perches).

The owl’s behavior was mostly restricted to flying between the two perches in a nearly straight line (Fig. 1B). Therefore, analyses and statistical tests in this study were reduced to 1D along the horizontal *x* direction (long axis of the room). Rare cases where owls landed on the ground or other positions in the room were excluded from analysis. To compute the firing-rate curves (1D firing-rate maps), we used a fixed spatial bin width of 10 cm. In each bin the overall number of spikes was divided by the bin’s occupancy (time spent), and the resultant firing-rate map was then smoothed with a 1D Gaussian kernel (σ = 1.5 bins). Firing-rate maps were calculated separately for each flight direction. Spatial rate maps in 2D (2D firing-rate maps: top view of XY plane; Fig. 1D) were used for display purposes only, and not for analyses. For producing 2D maps, spike-counts maps and occupancy maps were measured in 10×10 cm bins, 2D-smoothed with a fixed Gaussian kernel (σ = 1.5 bins), and then divided bin-by-bin to obtain the 2D firing rate map. This analysis included all bins occupied by the owl for at least 0.2 s (before smoothing).

To quantify the firing-rate differences between the two flight directions we calculated a discrimination index (DI):

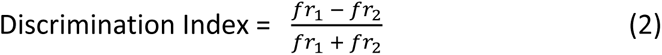

where fr_1_ is the average firing rate across all flights in one direction and fr_2_ is the same for the opposite direction. Cells that fired less than 20 spikes during flight were not included in this analysis. This index varies from –1 (firing only in one direction) to +1 (firing only in the other direction), with 0 indicating no directional discrimination. To assess the statistical significance of the discrimination index we used bootstrap statistics, as follows. The average firing rates of all individual flights in the session were intermixed and bootstrapped to produce a distribution of 1,000 discrimination indices. A result was considered significant if the real discrimination index was larger than 99% of the drawn indices (p<0.01).

To quantify the differences between the two perches or between flight versus perching, discrimination indices were calculated as described above. A similar bootstrap analysis was performed to assess significance of the discrimination indices. A value larger than 99% of the bootstrapped indices was considered significant.

Some of the owls showed a consistent behavioral preference to stand on one of the perches more than the other. Therefore, for perch discrimination analysis we equalized the time window for spike counts on both perches. If the owl stayed, for example, on the east perch for 15 seconds, then flew to the west perch and stayed there for 2 seconds, we used in both perches a time window of 2 seconds, starting after landing, to measure firing rates. This was repeated for all pairs of standing epochs (each east-perch standing epoch was paired with the subsequent west-perch standing epoch), taking the shorter time in each pair as the window for analysis in every pair.

To quantify the spatial modulation (place-tuning) of the firing rates along the flight, the 1D spatial tuning curve (1D firing-rate map) in-flight was smoothed with a 1D Gaussian kernel (σ = 1.5 bins, using 10-cm bins). The smoothed curve was then used to quantify the spatial-information index in bits/spike (Skaggs et al., 1996); however, we used a slightly modified spatial-information index, for the following reason. For calculating spatial information, the mean firing rate per bin (rate map) is classically multiplied by the occupancy probability (the relative time spent in each bin; Skaggs et al., 1996). However, the owls’ flights between the two perches followed a bell-shaped velocity curve, so the occupancy probability per bin was in all cases biased towards the edges (Fig. S6A). This implies that in our experiments, cells with place fields at the center of the path will provide less spatial information as compared with cells that had an identical field-size but were close to the edges of the path (close to the east/west walls). We therefore compensated for this bias by flattening the occupancy probability, i.e. rather than multiplying the spike-count by the probability per bin, as in (Skaggs et al., 1996), we multiplied by the average probability per bin – and hence, we call this index “spatial information under uniform occupancy assumption” (SI_u_):

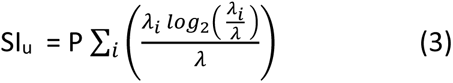

Where, λ_i_ is the firing rate in the i_th_ bin, λ is the overall average firing rate, and P is the average occupancy probability per bin (1 over the number of bins). For cells to be included in this analysis we required at least 20 spikes during flight in that direction.

To assess the significance of place-tuning during flight, we generated shuffled data sets, separately for the two flight-directions. Each spike train of each flight in a particular direction was randomly circularly shifted within the flight duration, thus generating a new set of spikes where the spike-train was maintained but the relation between spiking and position was broken; we then computed the firing-rate map and SI_u_ for each such shuffle. We repeated this shuffling 1,000 times, and thus 1,000 shuffled spatial-information indices were generated for each neuron. Cells for which the real SI_u_ exceeded the 99’th percentile of the corresponding shuffled distribution were considered as exhibiting significant place-tuning (p<0.01). To eliminate inclusion of cells with low spatial information but which were nevertheless significantly modulated due to a high firing rate, we imposed an additional criterion: an SI_u_ > 0.3 in at least one flight direction was required for placed-tuned cells.

To quantify the symmetry of the tuning curves between the two opposite directions we calculated a symmetry index (SymmI) as follows:

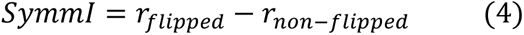

Where r*_flipped_* is the Pearson correlation coefficient between the firing-rate map in one direction and the corresponding firing-rate map in the other direction, but flipped in the east-west direction. r*_non-flipped_* is the Pearson correlation coefficient between the rate maps in the two directions, not flipped. If the pattern of the rate map displays a mirror-image, r_flipped_ will be close to +1 and _rnon-flipped_ will be closed to –1. Thus, this index varies between –2 to +2. Large positive values indicate symmetry (mirror image) – i.e. coding of distance, whereas large negative values indicate cells which code a similar place in both directions – i.e. coding of allocentric position.

### Histology

In 2 owls an electrical lesion was performed by injecting a positive current through one of the tetrodes (+5 µA for 20 sec). A week later, the owl was deeply anesthetized and perfused with phosphate buffer saline (PBS) solution, followed by 4% paraformaldehyde. The brain was removed and stored in 4% paraformaldehyde for 2–3 days at 4°C, then transferred to PBS. Following fixation, the owl’s brains were dehydrated in 70%, 80%, 95% and 100% ethanol, cleared in Xylene and embedded in paraffin wax. The paraffin-embedded brains were sectioned in the coronal plane at 5 µm using a microtome (Leica RM 2265). Sections were collected at 40 μm intervals, mounted on super-frost glass slides and dried in an oven at 37°C for 24 hours. After drying the sections, they were deparaffinized in xylene, rehydrated in a diluted ethanol series, and stained with 0.1% Cresyl violet solution (Nissl stain). The sections were then dehydrated, cleared and cover-slipped with DPX mounting medium (Merck). This allowed us to assign one recording location to HA_p_ (Owl TLV) and the other location to Hp (Owl DK). Recording locations in the other owls were assigned based on stereotaxic coordinates.

## Acknowledgments

We thank Tidhar Lev-Ari, Hadar Beeri, Liora Las and Shaked Ron for assistance and advice. This work was supported by research grants from the Rappaport Institute for Biomedical Research, the Adelis Foundation, and the Israel Science Foundation (grant no. 2655/18). Yoram Gutfreund also acknowledges the generous support of the Edward S. Mueller Eye Research Fund and the Irving and Branna Sisenwein Fund.

## Notes

### Competing Interest Statement

The authors have declared no competing interest.

