## Supplemental Material for "Spatial coding in the hippocampus of flying owls"

| Experiment # (Tetrodes implantation) | Owl Name | Brain location | # Sessions | # Single Units | Place tuning during West flights | Place tuning During East flights | Flight West pref. Cells | Flight East pref. Cells | Perch West pref. cells | Perch East pref. cells |
|---|---|---|---|---|---|---|---|---|---|---|
| 1 | Owl Arya | Left Hp | 40 | 220 | 44 | 29 | 38 | 13 | 31 | 36 |
| 2 | Owl Arya | Right Hp | 4 | 7 | 0 | 0 | 0 | 0 | 0 | 1 |
| 3 | Owl DK | Left Hp | 8 | 12 | 4 | 1 | 4 | 2 | 7 | 0 |
| 4 | Owl DK | Right Hp | 4 | 14 | 2 | 1 | 0 | 7 | 1 | 7 |
| 5 | Owl DB2 | Right Hp | 16 | 39 | 15 | 12 | 13 | 3 | 13 | 16 |
| | **Total Hp** | | **63** | **292** | **65** | **43** | **56** | **25** | **52** | **60** |
| 6 | Owl DB | Left HA | 11 | 49 | 19 | 17 | 9 | 8 | 12 | 9 |
| 7 | Owl DB2 | Left HA | 23 | 152 | 47 | 45 | 30 | 34 | 40 | 30 |
| 8 | Owl TLV1 | Right HA | 22 | 175 | 50 | 61 | 46 | 33 | 33 | 43 |
| | **Total HA** | | **56** | **376** | **116** | **123** | **85** | **74** | **85** | **82** |
| 9 | Owl DB2 | Left Wulst | 9 | 47 | 4 | 5 | 17 | 3 | 0 | 0 |
| 10 | Owl BB | Right Wulst | 10 | 46 | 9 | 13 | 9 | 1 | 1 | 3 |
| | **Total Wulst** | | **19** | **93** | **13** | **18** | **26** | **4** | **1** | **3** |

**Table S1:** The location of recordings, the number of sessions, and the number of cells recorded in each of the experiments.

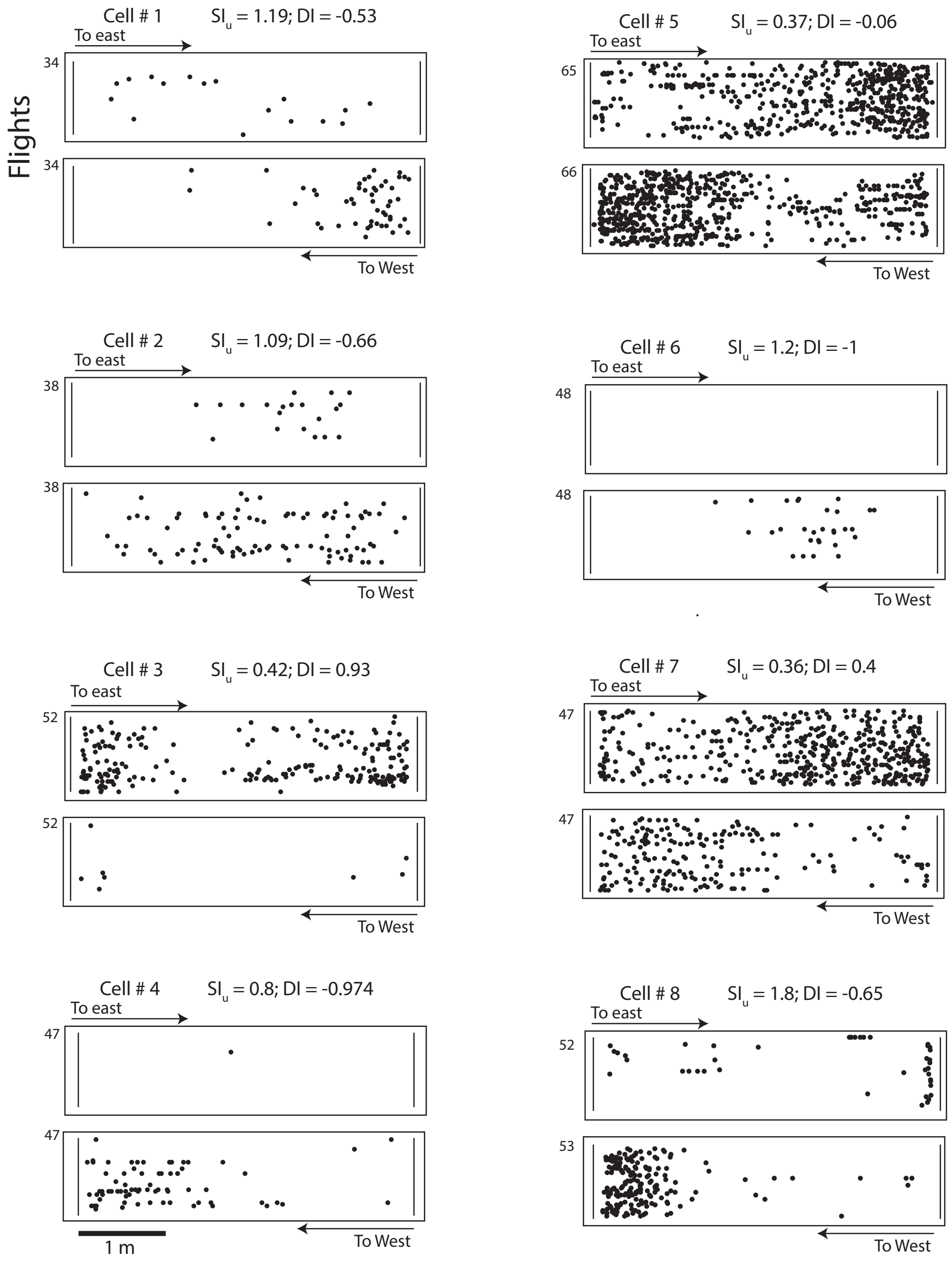


**Fig. S1.** Examples of 8 place - tuned cells recorded from the hippocampus (Hp), showing place tuning and directionality during flight. In each example, the upper raster shows flights to east and the lower raster flights to west. The number of flights in each direction is shown on the y-axis. The spatial information ($SI_u$) of the direction with the largest value and the discrimination index (DI) of flight direction are written above each example.

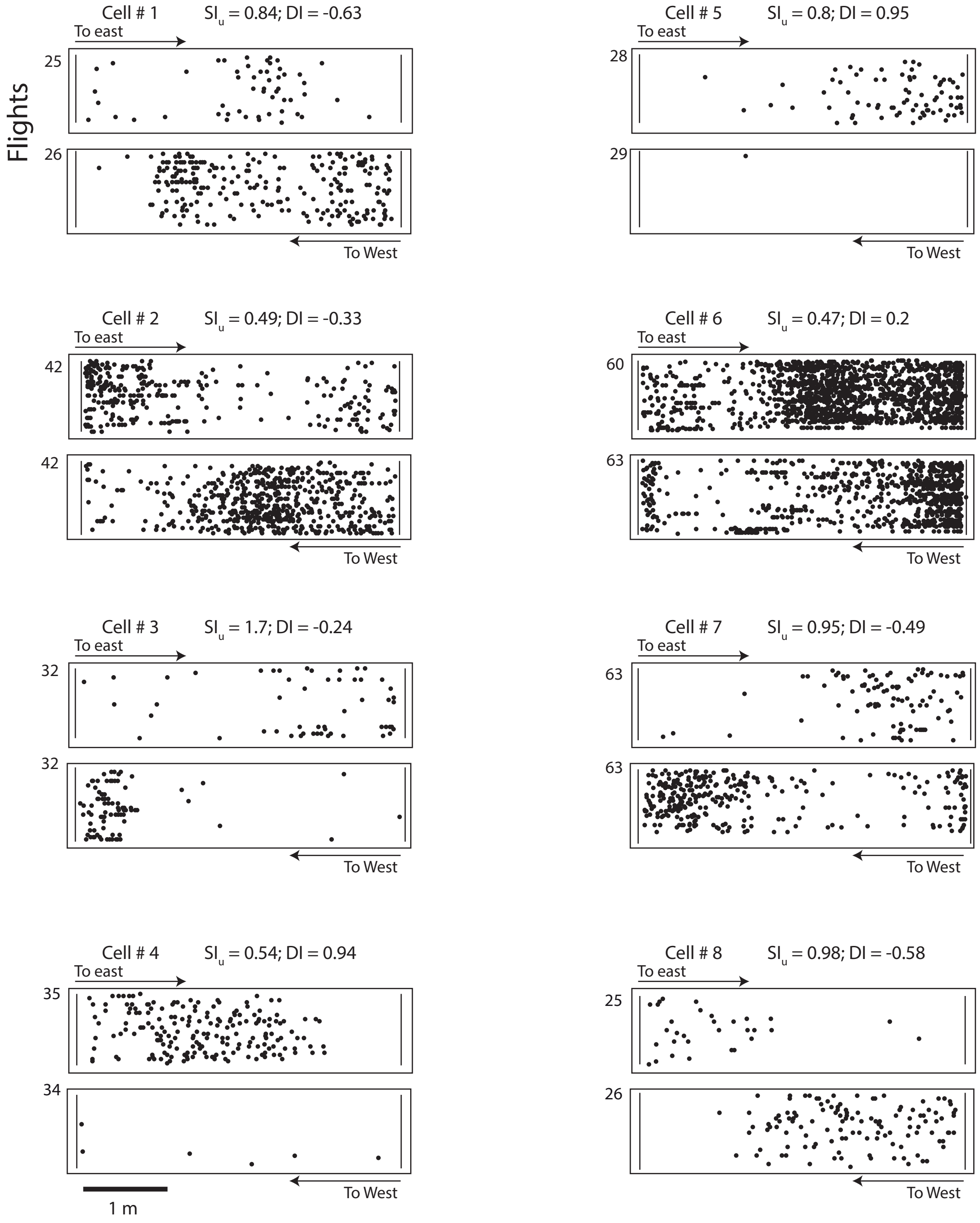


**Fig. S2.** Examples of 8 place - tuned cells recorded from the hyperpallium apicale posterior ($HA_p$). In each example, the upper raster shows flights to east and the lower raster flights to west. The number of flights in each direction is shown on the y-axis. The spatial information ($SI_u$) of the direction with the largest value and the discrimination index (DI) of flight direction are written above each example.

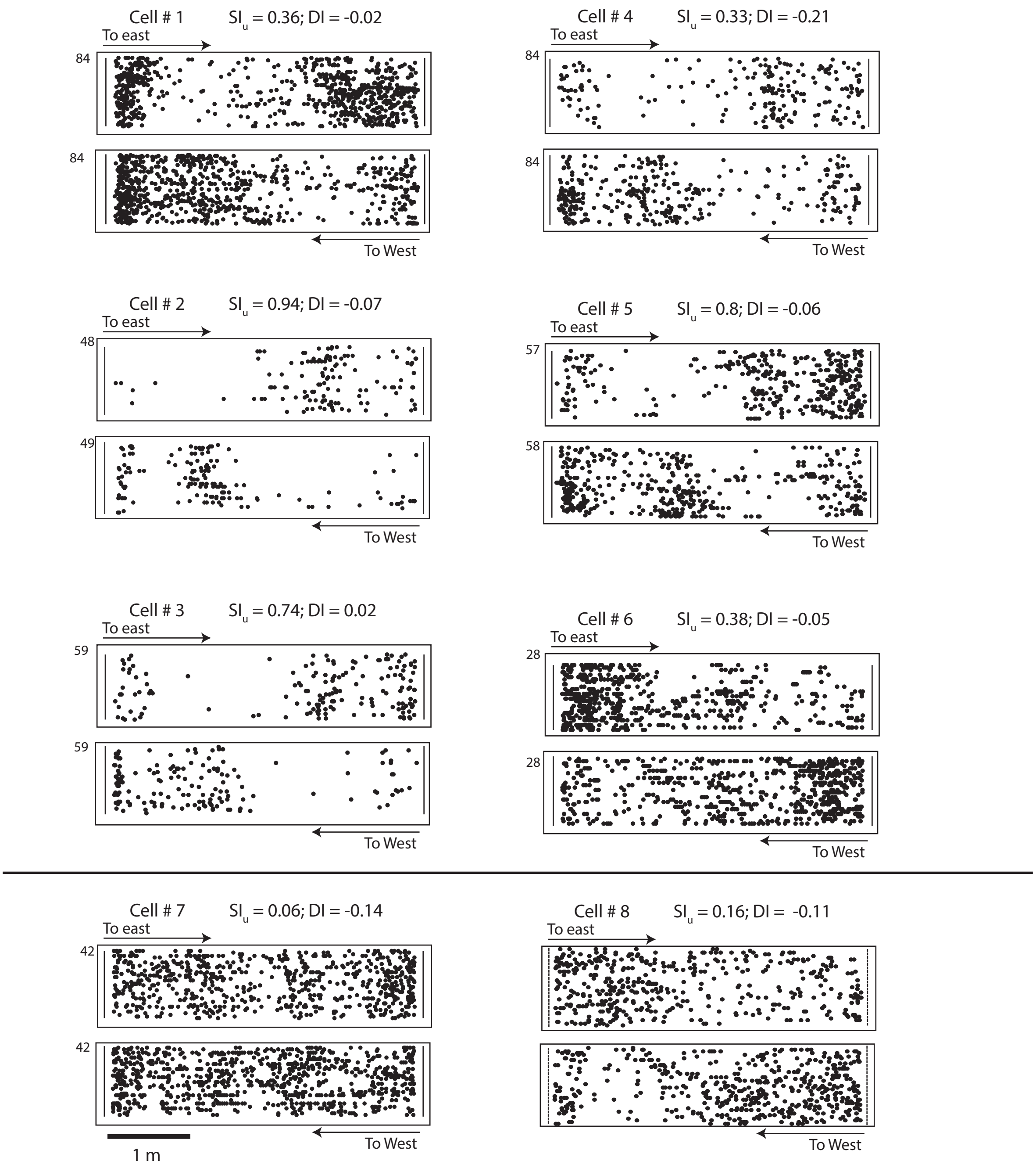


**Fig. S3.** Examples of 8 cells recorded from visual Wulst. Cells shown above the horizontal line are place-tuned ($SI_u > 0.3$). The number of flights in each direction is shown on the y-axis. The spatial information ($SI_u$) of the direction with the largest value and the discrimination index (DI) of flight direction are written above each example.

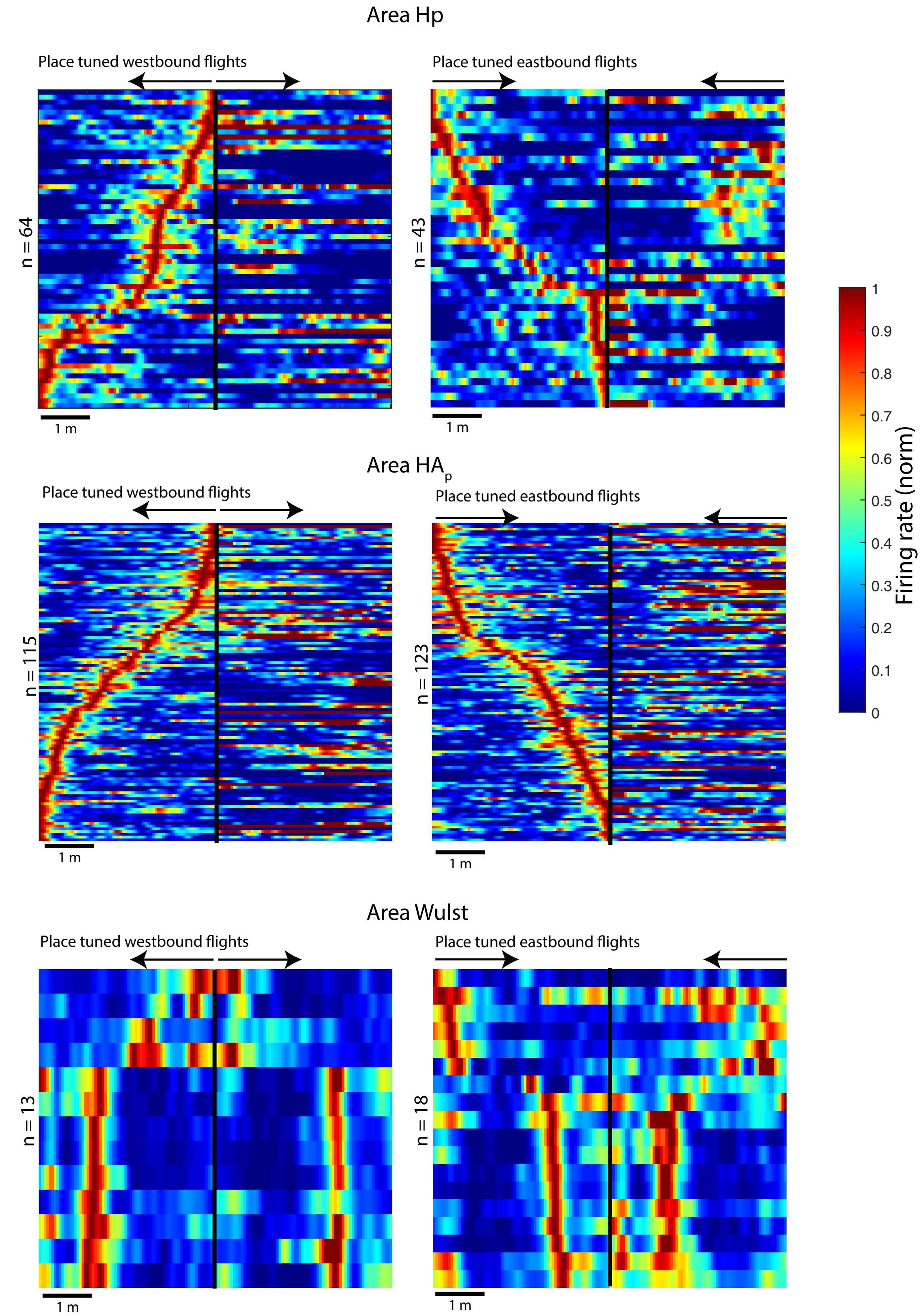


**Fig. S4:** Symmetry of responses in visual Wulst. Left column: All cells which showed place-tuned tuning curves for westbound flights are shown and sorted according to the position of peak firing from the onset of flight. Each line shows the normalized tuning curve of one cell in both directions. The black vertical line seperates westbound directions (significantly tuned and sorted) from eastbound directions of the same cells. The arrows show the onset and direction of flight. Right column shows the same, but for all cells whose tuning curves of eastbound flights were spatially-tuned. Results are seperated to Hp, HA and visual Wulst. In area Wulst, a clear mirror image appears in the corresponding opposite direction suggesting that Wulst neurons fire as a function of distance from take-off.

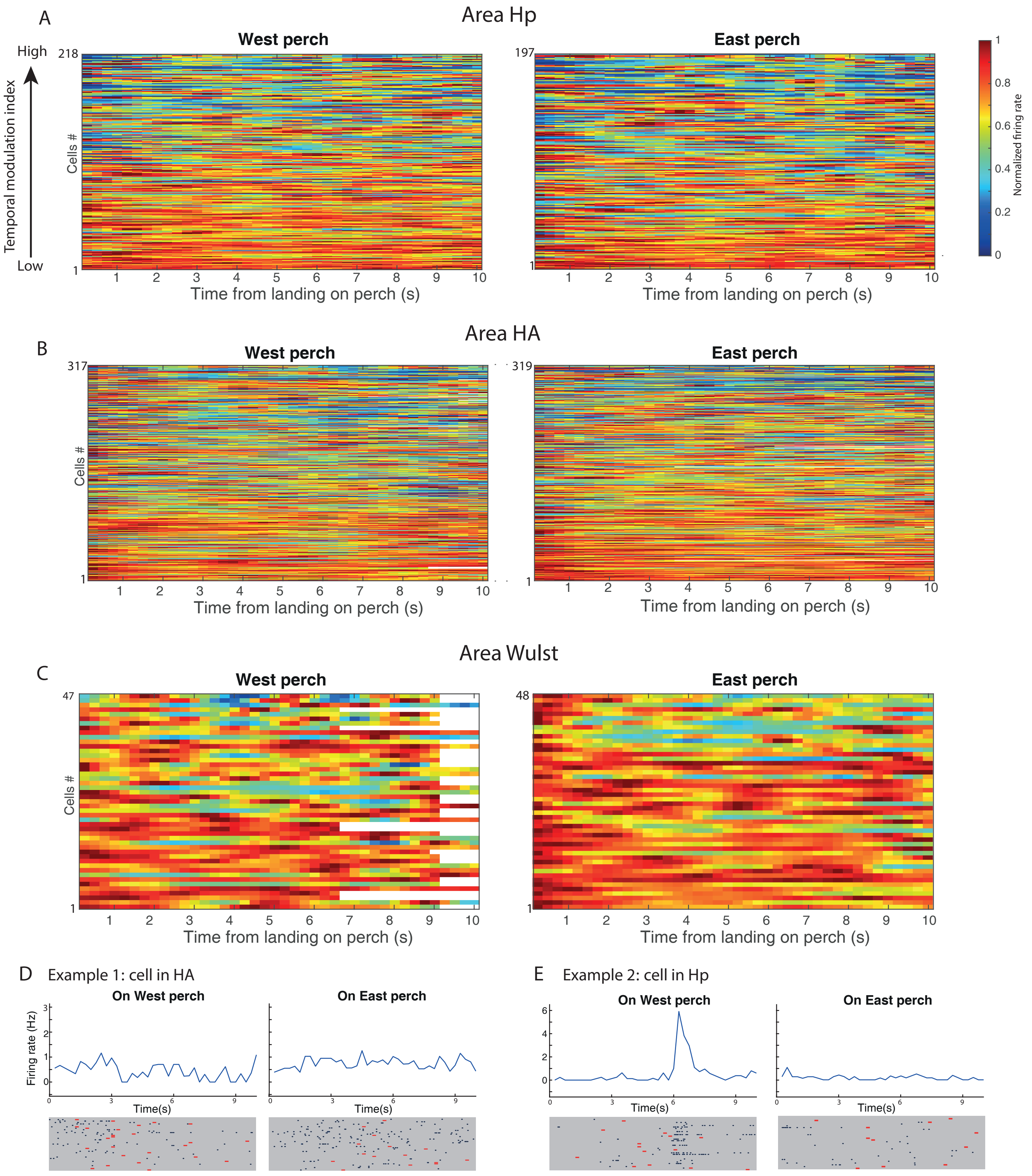


**Fig. S5.** Firing rates as a function of the time after landing. **A.** The smoothed and normalized firing rate curves in the 10 seconds following landing are shown for the neurons in Hp. On the left are results from the West perch and on the right results from the East perch. Neurons for which less than 50 spikes were fired over all, during the 10 seconds after landings were removed from the analysis. The first 500 ms after landing were removed from the analysis. Cells are sorted according to the temporal modulation index: $\mathrm{Sum}((I_i/I)\log(I_i/I))$ were $I_i$ is the firing rate at bin i and I is the average firing rate across all time bins. White pixels designate times for which the owl stayed less than three trials on the perch (departed the perch earlier than 10 seconds). **B** and **C.** same as in A but showing results for cells in the HA and area Wulst respectively. **D.** Exmaple of the firing rate curves and the corresponding spike rasters of a single cell in HA. Dots designate times of spikes, rows designate different trials and red bars designate the time of leaving the perch in the trials for which the owl stayed less than 10 seconds on the perch. **E.** Example of spike rasters and rate curves of a "time cell" recorded in the Hp. This was the only cell in our data set that showed such clear time coding. Layout of panel as in panel D.

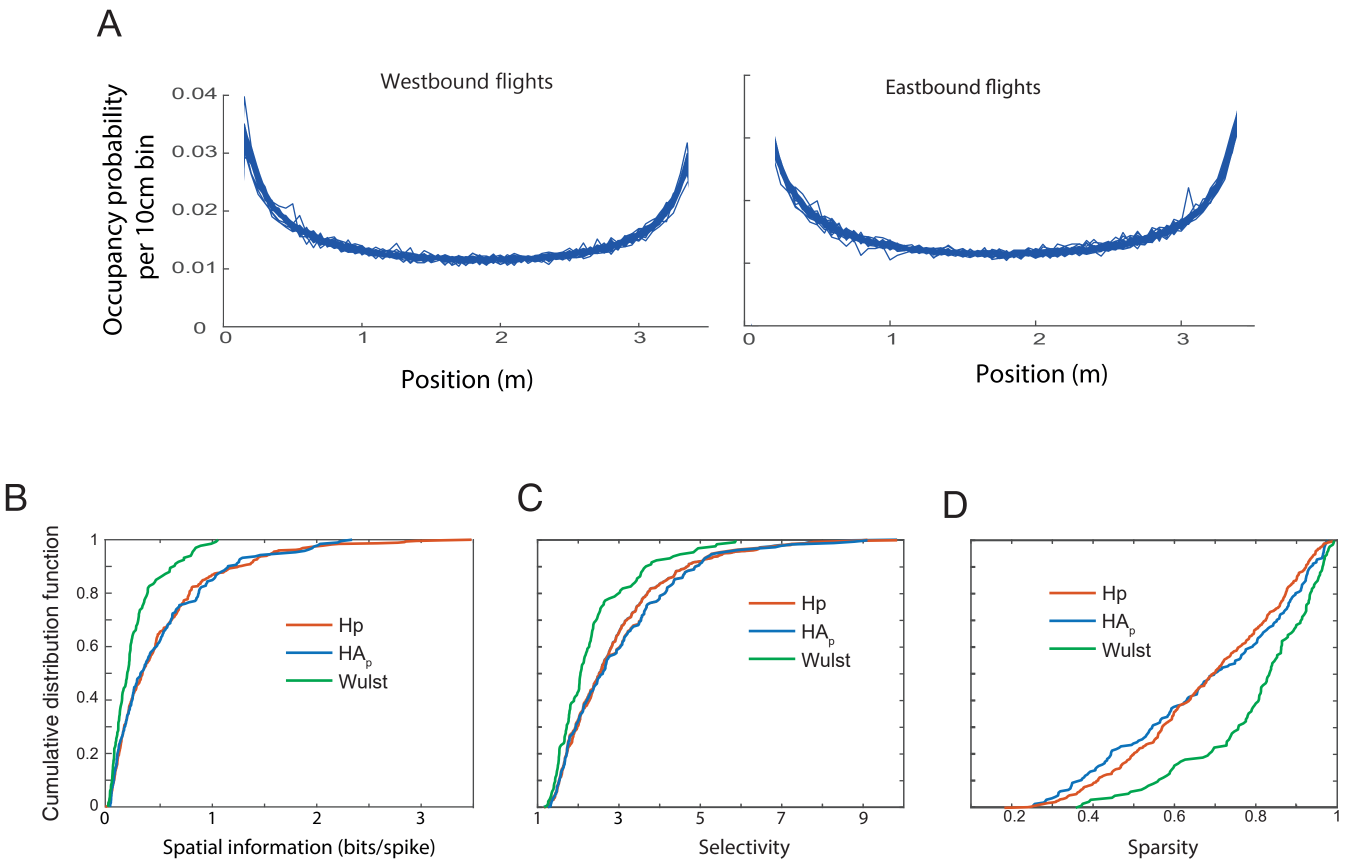


**Fig. S6: A,** Occupancy probability curves (the relative time spent at each 10 cm bin) from 40 different behavioral sessions of the same owl. The curves for westbound flights are on the left and for eastbound flights on the right. The occupancy curves are remarkably similar across flights and directions, following a bell shaped velocity curve. **B-D,** Comparison between the different brain regions of the cumulative distribution functions of three common spatial metrics: **B,** Spatial information:

$$\sum_{i=1}^{N} p_i \frac{fr_i}{fr} log \frac{fr_i}{fr}$$

Where N is the number of bins, $P_i$ is the occupancy probablity of bin i, *fr* is the mean firing rate and $fr_i$ is the firing rate of bin i. **C,** Selectivity: the maximum firing rate divided by the mean firing rate. **D,** Sparsity:

$$\frac{(\sum p_i fr_i)^2}{\sum p_i {fr_i}^2}$$

In all three metrics the spatial representations are significantly narrower in Hp and in $HA_p$ compared to area Wulst (Kolmogorov Smirnov test, $p<0.001$).
